# Comprehensive Guide of Epigenetics and Transcriptomics Data Quality Control

**DOI:** 10.1101/2024.08.02.606411

**Authors:** Arianna Comendul, Frederique Ruf-zamojski, Colby T. Ford, Pankaj Agarwal, Elena Zaslavsky, German Nudelman, Manoj Hariharan, Aliza Rubenstein, Hanna Pincas, Venugopalan D. Nair, Adam M. Michaleas, Stuart C. Sealfon, Christopher W. Woods, Kajal T. Claypool, Rafael Jaimes

## Abstract

Host response to environmental exposures such as pathogens and chemicals can cause modifications to the epigenome and transcriptome. Analysis of these modifications can reveal signatures with regards to the agent and timing of exposure. Exhaustive interrogation of the cascade of the epigenome and transcriptome requires analysis of disparate datasets from multiple assay types, often at single cell resolution, from the same biospecimen. Improved signature discovery has been enabled by advancements in assaying techniques to detect RNA expression, DNA base modifications, histone modifications, and chromatin accessibility. However, there remains a paucity of rigorous quality control standards of those datasets that reflect quality assurance of the underlying assay. This guide outlines a comprehensive suite of metrics that can be used to ensure quality from 11 different epigenetics and transcriptomics assays. Recommendations on mitigation approaches to address failed metrics and poor quality data are provided. The workflow consists of assessing dataset quality and reiterating benchwork protocols for improved results to generate accurate exposure signatures.

## Introduction

An epigenome consists of the current state of the chemical modifications to DNA and to the histone proteins that determine the packaging of the DNA. The main epigenetic modification to DNA is the methylation of cytosine to 5-methylcytosine which almost exclusively occurs on the cytosine of a CpG dinucleotide (i.e. when a cytosine follows a guanine). DNA methylation can inhibit the binding of transcription factors that can initiate transcription or by recruiting proteins involved in gene repression. (1) Another main epigenetic modification is the post-translational modification (PTM) of histones, commonly the acetylation of lysine residues. PTMs to histones are able to dictate the accessibility of DNA to transcription initiating proteins by influencing the association of positively charged histones to negatively charged DNA. The addition of a negatively charged functional group (i.e. acetyl) neutralizes the histone’s positive charge and weakens the protein’s association with DNA opening up access for transcriptional machinery to bind to regulatory regions and initiate transcription. (2) The transcriptome represents all coding and non-coding cellular ribonucleic acid (RNA) transcripts. Transcriptional regulation is often controlled by non-coding RNA (ncRNA), whereby ncRNA transcripts will bind to complementary messenger RNA (mRNA) to repress expression (3).

Some of these epigenetic and transcriptomic alterations can be transitory while other changes can persist for years to decades (4). Epigenetic exposure modifications, if distinctive, may provide potential patterns that can be combined together into exposure signatures (5–8). These epigenetic signatures may be gene and cell-type specific (5–8). Determining if an exposure signature is distinct can be difficult as exposures may only imprint subtle changes to the epigenome or transcriptome. These changes can be further masked if samples are analyzed in a pool containing poor quality samples. Without in depth quality control (QC), features extracted from a dataset may be the result of experimental conditions (e.g. batch effects) in the sample preparation process rather than the result of an exposure. For example, if partial sample degradation occurs at any point of the sample workflow, amplification for library construction will only occur on the non-degraded sequences resulting in a false representation of the original sample. Any sample processing step can leave compounding effects due to the potential of PCR amplification bias: the varying efficiency with which PCR amplifies sequences causing the end product after amplification to not accurately reflect the input DNA. Amplification bias often presents as the preferential amplification of sequences with high GC content and would be further exacerbated after any sample degradation. (9). As library construction for most epigenomic and transcriptomic sequencing requires PCR, just slightly mishandled sample prep steps can exponentially skew sequencing results. Because QC metrics are a result of the characterization of a sequence, they can hold insights into which sample preparation steps are potentially introducing these biases. Thus, leveraging QC metrics is a powerful tool to develop accurate and reproducible sample preparation protocols for epigenomic and transcriptomic datasets.

QC of epigenomics and transcriptomics datasets is often an overlooked step. Common bioinformatics workflows include bulked preprocessing steps for datasets that remove outliers and low-quality reads based on there deviation from all samples of a dataset opposed to QC assessment at the sample level prior to downstream analysis. Omitting poor quality samples prior to downstream analysis is essential to distinguish truly expressed genes from artifacts, variations introduced by non-biological processes. If a dataset is analyzed with a mix of poor and high quality samples, aberrant or differential gene expression could be removed as an outlier in the bulk preprocessing step, resulting in the loss or weakening of the signature (10). QC sample assessment at the individual level can also be used to determine if a subset of failed samples is grouped in a way that could indicate the source of the poor quality, like sharing the same sampling day.

In this article, QC pipelines that were established for 11 epigenetics and transcriptomics assays that calculate QC metrics for individual samples of a dataset were assessed. Herein, we review the potential cause for when reported metrics fall outside passing parameters and propose the best courses of mitigation.

The 11 assays assessed include Assay for Transposase-Accessible Chromatin using sequencing (ATAC-seq) (11), single cell ATAC-seq (scATAC-seq) (12), ChIPmentation (13), Infinium 850K MethylationEPIC BeadChip (14), Methylated DNA ImmunoPrecipitation sequencing (MeDIP-seq) (15), Multiplexed indexed T7 (Mint) chromatin immunoprecipitation (ChIP) sequencing (Mint-ChIP-seq) (16), micro ribonucleic acid (RNA) sequencing (miRNA-seq) (17), 10X Genomics chromium single cell multi-ome™(scMultiome) scATAC-seq and single cell RNA sequencing (scRNA-seq), RNA sequencing (RNA-seq) (18), scRNA-seq (19) or singular nuclear RNA-seq (snRNA-seq), and single nucleus methylcytosine sequencing (snmC-seq) and (snmC-seq2) (20, 21), see Figure 1.

**Fig. 1.**
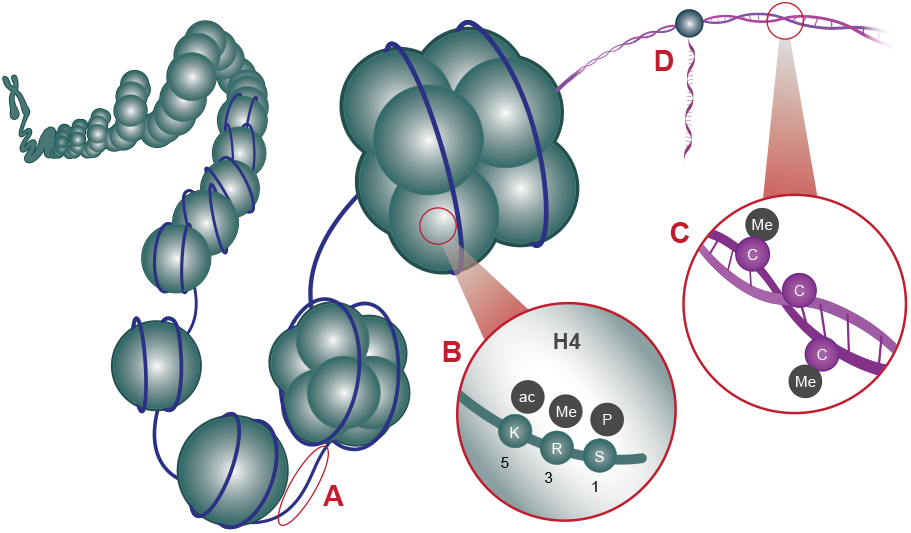
Epigenetics and transcriptomics assays/methods assess different aspects of gene regulation and expression, respectively. (**A**) Chromatin accessibility is measured by ATAC-seq and scATAC-seq (sometimes referred to as snATAC-seq for single nucleus). (**B**) Histone modifications are measured by MINT-ChIP-seq or ChIPmentation. (**C**) Cytosine (C) base methylation assessed by snmC-seq2, MeDIP-seq, or EPIC. (**D**) RNA/transcript sequencing by RNA-seq, scRNA-seq, or miRNA-seq. scMultiome assesses both scRNA-seq and scATAC-seq from the same cells.

## Methods

### Quality Control Metrics

Previously determined QC metrics were assessed for all assays at a good/pass/fail categorization. (22) These metrics cover a variety of attributes relating to sample quality such as sequencing depth, fraction of uniquely mapped reads, percent of aligned reads, etc. (Table 9). Single cell assays include further granularity in QC, including metrics such as number of cells and median unique molecular identifiers (UMI) per cell. For all assays, careful consideration is given to metrics that can point to biases potentially introduced from PCR steps.

### RNA-seq

The RNA content in both prokaryotic and eukaryotic cell consists of 80–90% rRNA, 10–15% transfer RNA (tRNA) and 3–7% messenger RNA (mRNA) and regulatory ncRNA. (23) RNA-sequencing (RNA-seq) is a nextgeneration sequencing (NGS) method that aims to quantify the mRNA transcripts in all cells from a given sample. Quantifying the transcriptome can reveal splice variants/isoforms, differentially expressed genes, novel transcripts and gene fusions that can characterize phenotypic variation in both disease etiology and epigenetic responses (24). When harmonized with epigenomic analysis, transcriptomic characterization can validate proposed epigenetic effects.

For RNA-seq, RNA was extracted from homogenized tissue or isolated cells (1,000 to 100,000 cells) (Direct-zol RNA MiniPrep kit). RNA concentration was assessed with fluorimetry using RiboGreen and RNA quality/integrity (RIN) and by capillary gel electrophoresis using a Fragment Analyzer. At this stage, poor-quality RNA samples, which either have a concentration below 8 ng/ul or a RIN below 7, are excluded as they may cause artifacts. If RIN is below 7, repeating the RNA extraction may be necessary. For samples meeting RNA quality requirements, complimentary DNA (cDNA) libraries were prepared using the universal Plus mRNA-Seq with NuQuant kit (Tecan Genomics) from poly(A) selected RNA. Sequencing library construction involved reverse transcription to generate cDNA with random and poly(T) primers followed by the addition of sequencing adaptors to each end of the cDNA fragment (25). Library concentration and the quality and extent of RNA degradation were evaluated by fluorimetry using Qubit and by capillary gel electrophoresis using Fragment Analyzer, respectively. Initially, shallow sequencing of the pooled libraries using a Mi-Seq system (Illumina) allows the confirmation of library quality and adjustment of the pools for deep sequencing. Deep sequencing of the libraries for up to 30 million reads per sample was performed on a Novaseq sequencer (Illumina).

QC checks are applied to ensure data quality that involve the analysis of raw reads, read alignment, transcript quantification and data reproducibility (26). Low sequencing depth may be from low sample input. Low uniquely mapped reads (i.e. reads that align directly to only one motif of the reference genome) below 75% can be the result of low library diversity. Low library diversity may be from low sample input to cDNA library construction or from PCR amplification bias. Optimizing PCR conditions for cDNA library construction by reducing cycles or tweaking heat steps can improve library diversity and can help achieve an optimal end GC content%. Integration of additional kits to workflows can be considered for sub-optimal ribosomal RNA(rRNA)% or %globinRNA. As rRNA is the predominant form of RNA, proper isolation of mRNA must be achieved for accurate transcript quantification. rRNA removal can enable higher sequencing depth of mRNA, leading to better detection of mRNA transcripts as well as uniquely mapped reads. (27) Globin-depletion prior to sequencing has been shown to improve the quality of RNA-seq data. (28) (29) Proper isolation of human peripheral blood mononuclear cells (PBMCs) (i.e. lymphocytes, monocytes, natural killer cells (NK cells) or dendritic cells) is essential to capture the RNA expression variations relevant to diseases and conditions in different cell types. Haemoglobin (Hgb) RNAs originate from red blood cells (RBCs) which can make up to 45% of the total volume of blood and 99% of the cellular volume. Thus, globin mRNA can compromise the detection of PBMC mRNAs by reducing the sensitivity to detect lower-level transcripts in the blood transcriptome. (30)

#### Quality Distribution

In addition to QC as part of the amplification and sequencing processes, evaluation of the outputs of any secondary analysis pipelines is imperative for understanding the quality of the algorithmic processing around assembly, alignment, quantifications, and any variant calling steps.

One example of this in RNA-seq and WES is the use of Picard^1^ (31), a tool from the Broad Institute, that enables the manipulation of high-throughput sequencing and contains functions to derive metrics around the quality of your alignment. Using functions such as Picard’s *CollectAlignmentSummaryMetrics, CollectBaseDistributionByCycle, CollectRnaSeqMetrics, CollectHsMetrics, CollectInsertSizeMetrics, MeanQualityByCycle, QualityScoreDistribution*, we can evaluate the quality of the sequencing inputs, but also the alignment performance of the sample. This is important in the assessment of the bioinformatics process that occurs after the assembly of reads through the secondary analysis steps.

The quality distribution metrics from Picard’s *QualityScoreDistribution* function, provides a count of reads by Phred quality score. This allows for the identification of quality issues where there may be significant proportion of lowquality reads that will affect subsequent analyses. Phred quality scores represent base-calling error probabilities using the equation:

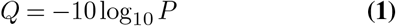

where *P* is the base-calling error probability and *Q* is the resulting Phred score. For example, a base-calling accuracy of 99% (or 1/100 error probability) would be represented with a Phred score of 20.

In Table 1 below, this example shows that 3.4% of the reads have a quality score of 11, which may be problematic, depending on the sensitivity of the research use case in question.

**Table 1.** Example of the quality distribution output from Picard’s *QualityScoreDistribution* function.

| QUALITY | COUNT_OF_Q |
| --- | --- |
| 11 | 457812700 |
| 25 | 708452935 |
| 37 | 12137945809 |

Furthermore, as part of the sequencing process, read quality should not vary significantly between cycles. Picard’s *MeanQualityByCycle* function also provides a breakdown of mean read quality by cycle. This view helps to identify if there is a significant change in read quality over time throughout the sequencing process. In Table 2 below, the example output shows that the mean remains stable (and high, > 20) through the first several cycles shown.

**Table 2.** Example of the mean quality output by cycle from Picard’s *MeanQualityByCycle* function.

| CYCLE | MEAN_QUALITY |
| --- | --- |
| 1 | 35.40155 |
| 2 | 35.58375 |
| 3 | 36.02574 |
| 4 | 36.13188 |
| 5 | 36.21469 |
| 6 | 36.53705 |
| 7 | 36.54161 |
| 8 | 36.57361 |
| ... | ... |

#### Alignment Metrics

Once the quality of sequencing reads has been deemed sufficient and an alignment has been performed, an additional QC step includes the evaluation of alignment performance. Picard’s *AlignmentSummaryMetrics, RnaSeqMetrics*, and *HsMetrics* functions provide convenient breakdowns of the reads and their alignment.

*AlignmentSummaryMetrics* shows the total reads versus the number of reads that were aligned along with metrics such as mismatch rate, mean read length, and indel rate. It is useful to confirm that the total reads aligned is on par with the reported number of reads from the sequencing process. Plus, this output can help identify if there is a high number of bad cycles, an unacceptable error rate, or an unacceptably high number of reads there were not aligned in pairs.

In Table 3 above, note how the mismatch and error rates are quite low. Plus, the high quality (HQ) aligned reads is a significant majority of all aligned reads (85,211,166 / 102,759,046 ≈ 83%) and 100% of the reads are aligned in pairs. If there were issues with the alignment quality, some of these metrics may help indicate the source of the issue(s). Furthermore, investigating the distribution of the bases within the transcripts of an RNA-seq experiment, we can better understand the makeup of the sample that was processed through an RNA-seq secondary analysis pipeline. For example, Table 4 below, we can see that 0 reads were ignored and that there were a very small number of bases which belong to rRNA and 95.8% of the bases were mRNA (messenger RNA), which is desired. Also, given that 25% of the human genome is introns (32) and that, in theory, RNA-seq samples should contain minimal intronic regions of the genome after downstream analyses, we can adjudicate this notion by seeing that <4% of the bases in the sample shown above are intronic bases.

**Table 3.** Example of the alignment metrics from Picard’s *AlignmentSummaryMetrics* function.

| CATEGORY | FIRST_OF_PAIR | SECOND_OF_PAIR | PAIR |
| --- | --- | --- | --- |
| TOTAL_READS | 54929952 | 54929952 | 109859904 |
| PF_READS | 54929952 | 54929952 | 109859904 |
| PCT_PF_READS | 1 | 1 | 1 |
| PF_NOISE_READS | 0 | 0 | 0 |
| PF_READS_ALIGNED | 51379523 | 51379523 | 102759046 |
| PCT_PF_READS_ALIGNED | 0.91668 | 0.91668 | 0.91668 |
| PF_ALIGNED_BASES | 5128423636 | 5125843525 | 10254267161 |
| PF_HQ_ALIGNED_READS | 42605583 | 42605583 | 85211166 |
| PF_HQ_ALIGNED_BASES | 4256380666 | 4255199751 | 8511580417 |
| PF_HQ_ALIGNED_Q20_BASES | 4220745549 | 4199744152 | 8420489701 |
| PF_HQ_MEDIAN_MISMATCHES | 0 | 0 | 0 |
| PF_MISMATCH_RATE | 0.002022 | 0.002443 | 0.002232 |
| PF_HQ_ERROR_RATE | 0.001804 | 0.002239 | 0.002022 |
| PF_INDEL_RATE | 0.000146 | 0.000147 | 0.000146 |
| MEAN_READ_LENGTH | 98.212138 | 98.182183 | 98.19716 |
| READS_ALIGNED_IN_PAIRS | 51379523 | 51379523 | 102759046 |
| PCT_READS_ALIGNED_IN_PAIRS | 1 | 1 | 1 |
| PF_READS_IMPROPER_PAIRS | 0 | 0 | 0 |
| PCT_PF_READS_IMPROPER_PAIRS | 0 | 0 | 0 |
| BAD_CYCLES | 0 | 0 | 0 |
| STRAND_BALANCE | 0.473177 | 0.506847 | 0.490012 |
| PCT_CHIMERAS | 0.005451 | 0.005451 | 0.005451 |
| PCT_ADAPTER | 0 | 0.000001 | 0 |

**Table 4.** Example of the RNA-seq metrics from Picard’s *CollectRnaSeqMetrics* function.

| CATEGORY | METRIC |
| --- | --- |
| PF_BASES | 12103166420 |
| PF_ALIGNED_BASES | 11274633239 |
| RIBOSOMAL_BASES | 4910 |
| CODING_BASES | 9300635422 |
| UTR_BASES | 1499010311 |
| INTRONIC_BASES | 370755932 |
| INTERGENIC_BASES | 104226665 |
| IGNORED_READS | 0 |
| CORRECT_STRAND_READS | 8479813 |
| INCORRECT_STRAND_READS | 285381 |
| NUM_R1_TRANSCRIPT_STRAND_READS | 121327 |
| NUM_R2_TRANSCRIPT_STRAND_READS | 4059650 |
| NUM_UNEXPLAINED_READS | 220590 |
| PCT_R1_TRANSCRIPT_STRAND_READS | 0.029019 |
| PCT_R2_TRANSCRIPT_STRAND_READS | 0.970981 |
| PCT_RIBOSOMAL_BASES | 0 |
| PCT_CODING_BASES | 0.824917 |
| PCT_UTR_BASES | 0.132954 |
| PCT_INTRONIC_BASES | 0.032884 |
| PCT_INTERGENIC_BASES | 0.009244 |
| PCT_MRNA_BASES | 0.957871 |
| PCT_USABLE_BASES | 0.892299 |
| PCT_CORRECT_STRAND_READS | 0.967442 |
| MEDIAN_CV_COVERAGE | 0.919178 |
| MEDIAN_5PRIME_BIAS | 0.231197 |
| MEDIAN_3PRIME_BIAS | 0.286529 |
| MEDIAN_5PRIME_TO_3PRIME_BIAS | 0.392208 |

It is also useful to note the percentage of the bases in your sample that were able to be aligned. In the example above, 93% (11,274,633,239 / 12,103,166,420) of the bases were aligned and the percentage of usable bases in the sample was 89%.

These aforementioned Picard outputs are just some of many metrics that may help to assess the quality of the results from any bioinformatics secondary analyses. While the quality of the input sequences may be high, issues with alignment may persist due to sample impurity, off target amplification, and other factors.

### scRNA-seq

Single-cell RNA-seq (scRNA-seq) is used to quantify coding and noncoding RNA transcripts of individual cells. scRNA-seq reveals the transcriptional cellular heterogeneity that is masked by bulk RNA-seq methods. This delineation can refine an exposure-based signature to be cell or tissue type specific.

Coupling microfluidic and nanodroplet-based systems with NGS enables the genome-wide expression profiling of thousands of cells (33). The scRNA-seq samples were characterized with the 10x Chromium Single Cell 3 Gene Expression assay kit. This method uses a microfluidic device to encapsulate single cells in a droplet that holds reverse transcription reagents. In each droplet the cell is lysed and a microreaction of reverse transcription occurs on polyadenylated RNAs, creating cDNA with capture sequences, UMI (unique molecular identifier) to distinguish separate RNA transcripts, and a unique cell barcode causing all cDNAs from the same cell to have the same barcode. The emulsion is then broken and pooled cDNAs undergo PCR amplification. The quality of the amplified cDNA was assessed with microcapillary-based electrophoresis using a Bioanalyzer. If low-quality RNA is indicated at this point, the RNA may have been degraded and new cell preparation or nuclei extraction is likely the best solution. cDNA is used for library construction. If the Bioanalyzer traces of the sequencing libraries show multiple peaks, library preparation should be repeated. Adapter and sample indices are incorporated into libraries for next generation short read sequence compatibility.

The quality and viability of cells at the start of any singlecell experiment is critical to achieve good QC metrics downstream. Cryopreservation and sample preparation technique discrepancies can cause variation in sample integrity and input quantity. As RNA degradation is closely related to sample preservation technique, ensuring proper storage and extraction methods can mitigate initiating the sequencing process with low and poor-quality input samples. If performing whole cell scRNA-seq, cell viability is assessed and should preferentially be above 70%. If the starting material are nuclei, assessing nuclei quantity using a fluorescence based cell counter is recommended. (34–36).

High mitochondrial reads in cells can be an indicator of damaged or dying cells. This is because in stressed cellular environments mitochondria can release proteins to induce apoptosis. (37) This can be caused by either dissection of necrotic tissue or by the stress caused on cells throughout the sample prep process disassociating cells from the tissue sample which can introduce artifacts to the sample. Also, if a cell is lysed, cytoplasmic mRNA can leak out through a broken membrane resulting in only mitochondrial RNA to be conserved and in turn over amplified and sequenced. (38, 39) If samples have high amount of cells with mitochondrial reads cell isolation techniques should be reviewed for optimization. (40).

RNA degradation could be the culprit for any failed metric. Working on ice and working quickly will alleviate RNA degradation. RNA degradation prior to cDNA synthesis can result in low cDNA yield and can cause the total number of cells detected to be low. Low cDNA synthesis yields will in turn cause low inputs for cDNA library construction. Low input into library construction can increase the chances of amplification bias in library construction resulting in low diversity libraries which can also cause a low number of cells to be detected in sequencing. Partial sample degradation of the cDNA library can result in low UMI per cell. However, median UMI counts may be skewed low if a sample contains a large proportion of neutrophils and granulocytes as they have a relatively low RNA content and high levels of ribonucleases (RNases) which results in fewer sequencing reads per neutrophil and granulocyte cell. If a low median UMI is not due to poor sample quality, optimal neutrophil and granulocyte sequencing reads may be achieved by increasing the PCR cycles by two during cDNA amplification. Deeper sequencing may also improve QC metrics.

### Targeted Gene Expression

As mentioned above, the input cell-level quality is imperative to the quality of downstream analyses. For example, the 10X platform also includes a QC toolkit called Cell Ranger (41). Cell Ranger’s *count* function provides count-level metrics per sample of the scRNAseq output. In the example output in Table 5 below, note the comparatively low number of cells found in the fourth sample along with its low fraction of reads that were found in cells. This may indicate an issue in the quality or integrity of the cells found in that sample.

**Table 5.** Example Cell Ranger targeted gene expression counts output. Failed samples are highlighted in red with an asterisk (*).

| Sample | 30-1006397_202401.SC-C1 | 30-1006342_202401.SC-D1 | 30-1006401_202401.SC-A2 | 30-1006422_202401.SC-C2 | 30-1006425_202401.SC-B1 |
| --- | --- | --- | --- | --- | --- |
| Estimated Number of Cells | 7,520 | 15,316 | 10,752 | 677* | 15,002 |
| Mean Reads per Cell | 79,382 | 34,812 | 45,758 | 537,410 | 24,610 |
| Median Genes per Cell | 1,312 | 3,624 | 870 | 609 | 134 |
| Number of Reads | 612,744,323 | 521,800,171 | 514,464,287 | 372,162,063 | 376,655,791 |
| Valid Barcodes | 98.50% | 98.50% | 98.50% | 97.50% | 97.50% |
| Valid UMIs | 100.00% | 99.50% | 100.00% | 100.00% | 99.99% |
| Sequencing Saturation | 93.50% | 51.50%* | 91.50% | 97.50% | 97.50% |
| Q30 Bases in Barcode | 96.20% | 96.30% | 96.20% | 95.60% | 95.60% |
| Q30 Bases in RNA Read | 95.10% | 95.40% | 95.60% | 94.20% | 94.40% |
| ... | ... | ... | ... | ... | ... |
| Fraction of Reads in Cells | 81.66% | 93.60% | 71.82% | 18.10%* | 81.23% |
| Total Genes Detected | 14,393 | 12,152 | 17,219 | 17,794 | 16,560 |
| Median UMI Counts per Cell | 1,441 | 691 | 1,408 | 259 | 727 |

Furthermore, this table shows that the percentage of bases that are high quality (Q30 or above) is >90% across all the sample as is the validity of the samples’ barcodes (>90%). So, the issue for the problematic sample shown above is not in the quality of the sequencing, rather, the cells in that sample. This type of output may also show sample impurity if the total genes detected is lower than expected or the mean reads per cell is also quite low.

Common issues in scRNA sequencing may arise from issues in sample integrity. For example, cells may have apoptosed or have been inadvertently lysed as part of the sample preparation process. This issue may be detected when looking at the cell-level QC output from Cell Ranger or other tools.

For example, Table 6 below shows that the percentage of cells that are predicted to be dead are quite high in the fourth sample from the above example.

**Table 6.** Example Cell Ranger cell-level QC predicting the number and percentage of dead cells. Failed samples are highlighted in red with an asterisk (*)

| Sample | Pct. Predicted Dead | N Predicted Dead |
| --- | --- | --- |
| 30-1006397_202401.SC-C1 | 0.1% | 8 |
| 30-1006342_202401.SC-D1 | 2.6% | 398 |
| 30-1006401_202401.SC-A2 | 0.0% | 0 |
| 30-1006422_202401.SC-C2 | 68.9%* | 466 |
| 30-1006425_202401.SC-B1 | 12.7% | 1905 |

It is expected for some cells to be dead as part of any input sample. However, sequencing performed on older samples or those that have been frozen and thawed improperly may increase the quantity of dead cells. Then, when a sequencing process results in few viable cells and a low percentage of in-cell reads, the presence of a high percentage of dead cells further hinders the samples viability for downstream processing and usage.

### Cell Classification

A common goal of scRNA-seq may be to characterize the variation in gene expression by cell type across samples. For a given study, cells can be classified into a number different cell types based on an input gene signature matrix. For example, using an R library such as SingleR, cells can be mapped to a variety of cell types based on a their gene expression. In the example data shown in Table 7 below, there is varying quantities and percentages of the few cell types that are defined.

**Table 7.**
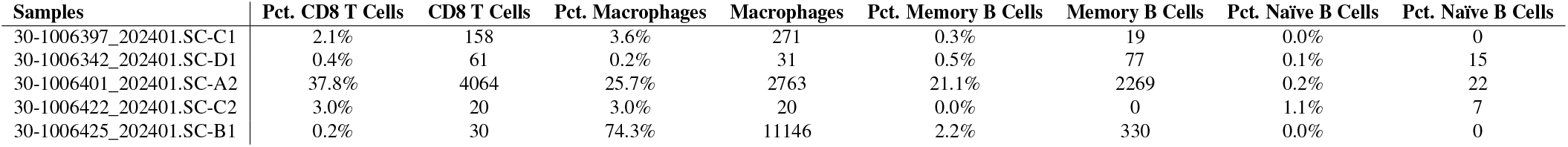
Example cell type mapping from a scRNA-seq analysis.

| Samples | Pct. CD8 T Cells | CD8 T Cells | Pct. Macrophages | Macrophages | Pct. Memory B Cells | Memory B Cells | Pct. Naïve B Cells | Pct. Naïve B Cells |
| --- | --- | --- | --- | --- | --- | --- | --- | --- |
| 30-1006397_202401.SC-C1 | 2.1% | 158 | 3.6% | 271 | 0.3% | 19 | 0.0% | 0 |
| 30-1006342_202401.SC-D1 | 0.4% | 61 | 0.2% | 31 | 0.5% | 77 | 0.1% | 15 |
| 30-1006401_202401.SC-A2 | 37.8% | 4064 | 25.7% | 2763 | 21.1% | 2269 | 0.2% | 22 |
| 30-1006422_202401.SC-C2 | 3.0% | 20 | 3.0% | 20 | 0.0% | 0 | 1.1% | 7 |
| 30-1006425_202401.SC-B1 | 0.2% | 30 | 74.3% | 11146 | 2.2% | 330 | 0.0% | 0 |

In the example above, the third sample seems to have a much larger ratio of memory B cells and CD8+ T cells as compared to the others. While this may not be problematic, it may be indicative that the samples are quite different. This is important in studies where the samples (that are perheps from different study participants) need to be of the same tissue types to get as close to a one-to-one comparison. In this example set, it becomes clear that the input tissues may not have been the same or the state of those tissues certainly varies.

Also, it is important to pre-define the cell types of interest as this changes the required reference input for the bioinformatics mapping process. In the example above, only 4 immunorelated cell types were used. However, it completely misses other cells that may have been of interest in the samples (e.g., erythrocytes, plasma cells, keratinocytes, etc.).

### ATAC-seq

Assay for Transposase-Accessible Chromatin sequencing (ATAC-seq) is a fast and high-throughput method that has been widely used to profile chromatin accessibility across the genome. (11, 42–44). ATAC-seq identifies open chromatin regions (OCRs) by sequencing the 20-90 base pair (bp) sequences that connect nucleosomes. These open chromatin regions are often regulatory elements like promoters, enhancers, insulators and silencers that transcription factors bind to to influence and control gene expression. (45).(46). ATAC-seq offers valuable insight as most RNA polymerase activity to initiate transcription co-occurs with OCRs. (47) By coupling the same biological sample with ATAC-seq and downstream sequencing applications, we inferred underlying gene regulatory mechanisms by correlating differential chromatin accessibility with differential gene expression.

Nuclei extraction was performed. Genomic DNA was exposed to Tn5, a highly active transposase that targets open chromatin sites, and simultaneously cuts the ends of open chromatin regions and ligates sequencing adaptors to them to create fragments that can be PCR-amplified. Library preparation was done with Nextera DNA Flex Library Prep. Library quantity was assessed by microcapillary-based electrophoresis using a Bioanalyzer. Amplified libraries were checked to exhibit a fragment size distribution characteristic to nucleosomal-free region (NFR) lengths. Fragment size distribution of the amplified libraries is expected to show a characteristic nucleosomal laddering pattern with a periodicity of 200 bp, corresponding to DNA fragments from nucleosome-free regions (NFRs). High quality libraries were then purified and subjected to paired-end sequencing on a Nova-Seq instrument.

If sequencing depth is below 25 million, repeating the transposase reaction and library prep may be required to achieve proper sequencing depth. Over 40 million non-duplicate reads is indicative of strong library complexity. A high duplicate rate may be an indicator of a low library complexity, for which increasing initial cell input, repeating library preparation, or adjusting the volumes of low concentration samples when making a new pool for sequencing may resolve the issue. Although ATAC-seq is intended to primarily sequence regions in between nucleosomes, a high quality ATAC-seq sample will also include a sequence peak for one nucleosomal sequence which enables proper alignment of the sequences.

Signal-to-noise metrics are essential to look at in ATACseq since highly accessible DNA is a characteristic for cells of poor viability like activated granulates or dead and dying cells. Key signal-to-noise metrics are FRiP (Fraction of Reads in Peaks) and TSS (Transcriptional start sites) enrichment scores. Peaks are determined based on a 10 to 30-fold enrichment compared to background. (48) High FRiP scores indicate the experiment has a high signal-to-noise ratio as a majority of the reads are in peaks, enriched regions compared to background. A low FRiP score indicates the experiment to have had a low specificity as a majority of the reads are in non-specific regions. TSS enrichment score is assessed as an aggregate distribution of reads centered on TSSs and extending to 2000 bp in either direction. TSS regions should be enriched as they are typically in accessible chromatin regions. To ensure strong signal-to-noise ratios cell viability and sample integrity must be ensured. Repeating the transposition step would be necessary if the sample indicated non-specific sequencing.

#### scATAC-seq & snATAC-seq

Single nucleus ATAC-seq (snATAC-seq) and single cell ATAC-seq (scATAC-seq) generate profiles of chromatin accessibility at a single cell resolution. (49). Further delineating the OCRs of a sample to single cell granularity enables a more comprehensive view of a sample’s regulatory landscape. (50).

Similar to ATAC-seq, for scATAC-seq nuclei are isolated and the genomics DNA is treated with a transposase to cut open chromatin sites and ligate barcodes to the both ends of the sequence. Nuclei are then redistributed using a cell sorter and lysed. Nuclei are then pooled and are introduced to a second barcode by PCR with indexed primers complementary to the transposase-introduced adapters. All PCR products are then pooled and sequences are further identified as certain cell types using known cell type marker genes.

The same mitigation steps for ATACseq from the same failed metrics can be taken for scATAC-seq. Short sequence length may be indicative of sample degradation. (51–53)

#### scMultiome

Single cell Multiome (scMultiome) sequencing pairs scATAC-seq and scRNA-seq to simultaneously generate a cell-type-specific chromatin accessibility and transcriptional profiles in the same cells. (54) These concurrently made profiles can identify the variations in chromatin accessibility for the regulatory regions of a certain gene that result in the same transcriptional phenotype. The ability to take multi-omic measurements in parallel allows scMultiome to provide unambiguous inferences on genetic heterogeneity. (55–57)

The 10x Genomics Chromium Next GEM Multiome ATAC GEX (Gene Expression) kit was used to generate singlecell libraries. First nuclei are isolated and are bulk treated with Tn5 to generate chromatin accessible fragments. A microfluidic chip is then used to partition nuclei into individual emulsion droplets where UMI barcodes are added to the transposed DNA and mRNA. Droplets are pooled for preamplification PCR to ensure maximum recovery of barcoded transposed DNA and cDNA fragments. This PCR product is then used as input for ATAC library construction and cDNA amplification for gene expression library construction. Libraries were sequenced on a NovaSeq instrument.

Entering the assay with high-quality nuclei is critical for assay success. Extracellular debris in the sample can clog the microfluidic channels that partition nuclei into emulsions which can result in a low number of cells detected by sequencing. Improper emulsification from debris can also increase the chances of doublet formation, when two or more cells are in one emulsion droplet resulting in transcripts from more than one cell receiving the same cellular barcode. Minimizing the chances of doublet formation is critical for transcripts to be mapped to the proper cell type to enable overall accurate cellular annotation. Low quality nuclei can also cause nonhomogeneous emulsification reactions which can result in low median UMI detected per cell. Low GEX and ATAC sequence lengths can be indications of sample degradation. QC of GEX from a scMultiome sample is visualized in Fig 2. High quality samples have dinstinct clustering indicative of different cell types that are identifiable by the distribution of cell specific marker genes. (58)

**Fig. 2.**
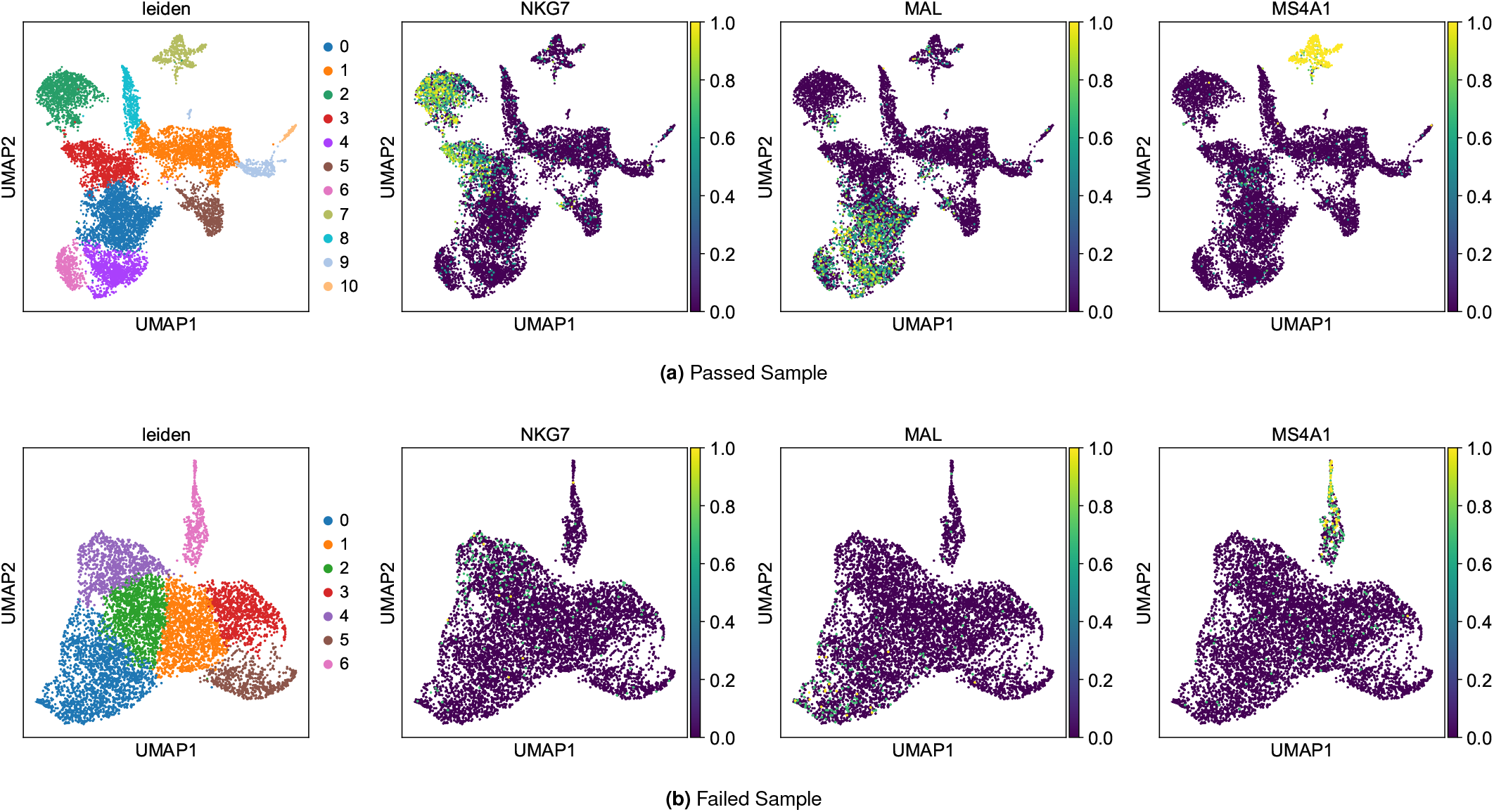
QC Visualization of scRNAseq data from PBMC samples from individuals exposed to pentaerythritol tetranitrate (PETN), a nitrate ester explosive. Each point represents a cell. Clustering is driven by transcriptomic profiles. Leiden clustering and distribution of marker genes for NK cells (NKG7), CD4 T cells (MAL) and B cells (MS4A1) shown for a passed sample (a) and failed sample (b).

### MethylationEPIC

DNA methylation is an epigenetic mark that is involved in regulating several cellular processes including gene transcription, genomic imprinting, and X-chromosome inactivation. DNA methylomes, the representation of methyl groups on a genome’s 5’-CpG-3’ dinucleotides sites added by methyltransferases, can serve as maps to environmental changes such as exposure to chemicals or pathogens to inform epigenetic signature development. (59) The cell-free methylome can also be determined by methylationEPIC analysis by inputted cell-free DNA (cfDNA) to detect abnormalities in methylation patterns indicative of cancer. (60) MethylationEPIC analysis via the Infinium MethylationEPIC v2.0 BeadChip provides quantitative genomewide CpG methylation screening at a single nucleotide resolution for over 850,000 CpG sites. (14) Probes are only designed to interrogate known CpG sites that have been demonstrated to be enhancers and regions associated with tumors. (61)

Input DNA first undergoes bisulfite conversion in a thermocycling process causing unmethylated cytosine bases to deaminate to uracil; methyl-cytosine bases remain as cytosine. Bisulfite converted DNA then underwent whole genome isothermal amplification and were enzymatically fragmented. DNA was purified and added to bead bound probes where the DNA hybridizes to the CpG locus specific oligomers on individual beads. Bead oligomers will either correspond to the CpG locus of the methylated (C) or the unmethylated (T) state. Next, allele-specific single base extension of the probes was conducted with either biotin labeled C and G nucletotides or dinitrophenyl labeled A and T nucleotides. Nucleotide labels are incorporated to improve signal-to-noise ratios. The array was fluorescently stained, scanned and the intensities of the unmethylated and methylated bead types at each locus were measured to determine the relative degree of locus methylation.

The EPIC array data quality checks include:

(i) 17 control metrics defined by the manufacturer (62), including Bisulfite Conversion II, which monitors successful bisulfite conversion Samples with a Bisulfite Conversion II metric below 1 might be partially converted, leading to inaccurate estimates of methylation levels. Also, samples with too many undetected probes or low overall fluorescence intensity are excluded. As fluorescence intensities of a SNP locus are calculated from two probes targeting either the wild type or the common mutant variant, the distribution of the proportion of methylated strands of a SNP locus (β-value) falls into three disjunct clusters: the heterozygous and the two homozygous genotypes. β-value deviation from the ideal trimodal distribution can serve as a means of outlier identification and a metric for poor technical performance. (62),
(ii) a sex check to detect mislabeled sex-discordant samples,
(iii) an identity check for fingerprinting sample donors,
(iv) a measure of sample contamination based on probes querying high-frequency SNPs.

Additionally, undetected probes should be filtered out from analysis to reduce spurious values (63).

Additional quality checks consist in evaluating sample clustering, background noise, probe variability, methylation call rate, beta value distribution, detection P-value distribution, and principal component analysis (PCA), as described below:

1. **Sample clustering:** The sample clustering should clearly separate samples into appropriate groups, with minimal overlap between groups.
2. **Background noise:** The background noise levels should be low and consistent across all arrays, with values less than 2% of the average signal intensity.
3. **Probes with high variability:** The proportion of probes with high variability (CV > 30%) should be low, typically less than 10%.
4. **Methylation call rate:** The methylation call rate should be high, with a minimum of 95% of probes with a valid methylation call in each sample.
5. **Beta value distribution:** The beta value represents the proportion of methylated strands for each CpG site. The beta value distribution should be within the expected range (0-1), with a minimum balance of probes with extreme values (less than 0 or greater than 1).
6. **Detection P-value distribution:** A low signal-to-noise ratio of fluorescence intensities results in probes with high detection p-values. Such probes should be removed as they are unreliable. The detection P-values should have a minimal proportion of probes with P-values greater than 0.05, typically less than 5%.
7. **PCA:** The first two principal components should explain a significant proportion of the sample variation (typically >50%), with minimal overlap between samples of different groups.

Input DNA quantity and quality influence assay performance. Failed probe detection can be the result of a high background. Lowering background may be alleviated by ensuring optimal amount of input DNA for the bisulfite conversion kit (being between 200-500 ng). High levels of background are also alleviated by optimizing PCR conditions in whole genome isothermal amplification - optimal primers and high annealing temperatures between 55-60 °C coupled with hot start polymerases can help since the template is AT -rich due to the non-methylated cytosine conversion to uracil in the PCR template. Proper reaction times for the initial bisulfite reaction could be also important; although methylated cytosine base deaminate at about two orders of magnitude slower overextending the bisulfite reaction time could lead to methylated cytosine to undergo the deamination yielding inaccurate results.

One drawback of MethylationEPIC is the background noise resulting from the off-target binding of probes due to their sequence homology. (64) Another source of noise in the data is the cross-talk between the green and red fluorophores used to differentiate the nucleotides that are incorporated in the single-base extension step. Beta value variations can occur depending on which instrument is used in quantifying signal intensities. [Illumina iScan, NextSeq 550, or NextSeq 550Dx systems.]

### Mint-ChIP-seq

Mint-ChIP-seq is a high-throughput method for genome-wide mapping of nucleosomes with specific histone modifications, such as H3K27ac and H3K4me3; H3K27ac is acetyl modified lysine residue of histone H3 and H3K4me3 is three methyl residues added to lysine residue 4 of histone H3. In this method, cross-linked chromatin is first pulled down using an antibody against the histone modification of interest and the corresponding genomic region is sequenced. (16) These histone modifications can alter chromatin structure and influence the accessibility of genomic regions to regulatory proteins and transcription factors. Histone specific signatures from Mint-ChIP have been used in characterizing disease mechanisms. (65)

Traditional ChIP-seq assays require a large number of input cells for sample preparation (66). The Mint-ChIP-seq uses a method that simultaneously fragments and digests chromatin using micrococcal Nuclease (MNase) from a small number of cells that have been optimized for minimizing sample loss and conducting chromatin Immuno-precipitation (IP). Double stranded barcoded adapters are ligated to the 5’ end of the fragmented nucleosome DNA. These adapters, referred to as Index #1, contain:

- T7 promoter
- Illumina SBS3 PCR priming sequence
- Barcode that is unique for each sample
- 5’ C3 spacer to prevent self-ligation

The T7 promoter is used for DNA amplification in a transcription reaction where multiple copies of RNA for each nucleosomal DNA are generated, which greatly reduces the amount of input cells required. Reverse transcription using primers that contain a PCR priming sequence on both ends generates cDNA that is PCR amplified for downstream library generation.

Throughput in ChIP-seq assays is low because each sample is processed individually. The Mint-ChIP assay yields a higher throughput by using a pool-and-split multiplexing approach where samples are pooled together using the sample barcode in the Index #1 adapter. The pooled samples are then split into individual ChIP assays, one for each histone modification mark and total H3 that serves as the control. Each split sample undergoes an immunoprecipitation reaction using an antibody specific to the histone mark or H3 histone (control) in a parallel reaction which increases the throughput (H3, H3K4me1, H3K4me3, H3K9me3, H3K27ac, H3K27me3, and H3K36me3).

A quantitative comparison of histone modification across samples in traditional ChIP-seq assay is difficult because separate IP reactions are required for the samples, and they are sensitive to the amount of starting material. Some strategies that have recently been used include incorporating exogenous DNA or synthetic histone spike-in control, but they don’t work well for small numbers of cells. The pool-and-split approach which allows for processing multiple samples in the same assay allows for quantitative normalization and comparison across samples.

After the samples are split, one for each histone mark, a second barcode, Index #2, is added which can identify the individual histone modification (ChIP assay). The libraries are then combined, which now contain different samples, represented by Index #1, and different histone marks, represented by Index #2, and sequenced using pair-end sequencing. The FASTQ files generated from the sequencing are then demultiplexed for each histone mark by Index #2 followed by demultiplexing for each sample using Index #1.

The Mint-ChIP-seq assay involves additional steps compared to the traditional ChIP-seq assay which has the potential to introduce bias. Quality control steps therefore become critical before Mint-ChIP data can be analyzed. As with most sequencing experiments, some QC parameters that should be checked for each sample and histone mark as an indication of a high-quality library include:

- Number of reads that uniquely map to the genome, which should be over 2 million but ideally greater than 3 million.
- Percentage of uniquely mapped reads that map to the genome (alignment percentage), which at minimum should be greater than 60 percent, but for a high-quality library is expected to be greater than 85 percent.

The key objective of a ChIP-seq assay is the identification of peaks, which are regions of the genome where the histone modification is expected, and is identified by the enrichment of alignment of reads in these regions. The presence of reads indicates the presence of the histone mark. FRIP is used to assess signal-to-noise ratio. Ideally all the aligned reads should fall within peaks but a value of greater than 40 percent is considered acceptable due to presence of background reads from regions other than the peaks. In traditional ChIP-seq assay the background signals are controlled using histone spike-in or by incorporating exogenous DNA. This approach is not appropriate for the Mint-ChIP assay due to low number of cells but is overcome by pooling samples in the same IP reaction and using total Histone (H3) as control.

ChIP-seq assays suffer from high PCR duplication rate due to the low amount of immunoprecipitated DNA which requires higher PCR amplification. In Mint-ChIP assays this is further exacerbated due to the two-step amplification which leads to higher PCR duplicate rate and low complexity, especially for histone marks that are not commonly found on the genome such as H3K27ac. The PCR duplicate rate for these marks is high and difficult to control. We defined a new parameter called useful reads efficiency, which could better capture the usefulness of the deduplicated reads. Useful reads efficiency is calculated by dividing useful reads by all reads. Useful reads are those that uniquely map to the genome after removing the PCR duplication. The useful reads efficiency should be greater than 70% for optimal quality. Typically if the PCR duplication rate is too high, the useful reads efficiency will decrease. Due to the expected variation in the PCR duplication rate, it is suggested to set a threshold based on specific marks. Recommend rates are listed in Table 8. A lower PCR duplication rate is better and leads to higher useful reads efficiency.

**Table 8.** Mint-ChIP suboptimal PCR duplication rate for specific histone marks.

| Histone Mark | PCR Duplication Rate (max) |
| --- | --- |
| H3K4me1 | 70% |
| H3K4me3 | 40% |
| H3K27me3 | 40% |
| H3K36me3 | 60% |
| H3K9me3 | 60% |
| H3 | 70% |
| H3K27ac | 40% |

**Table 9.** Quality metrics for different assays. Bounds are given for failing, passing, and excellent, with suggested mitigating procedures to improve quality. * Remove sources of sample degradation. ** Remove sources of sample contamination.

| Assay | Metric | Fail | Pass | Best | Mitigations |
| --- | --- | --- | --- | --- | --- |
| ATAC-seq | Sequencing depth | <25M | $\geq 25M$ | | *<br>Repeat library preparation |
| | %Aligned reads | <50% | [50%, 75%) | $\geq 75\%$ | Repeat nuclei extraction |
| | Non-duplicate reads | <15M | [15M, 40M) | $\geq 40M$ | Repeat library preparation<br>increase initial cell input<br>concentrate sample for<br>sequence input |
| | Overlap FRIP | <0.05 | [0.05, 0.1) | $\geq 0.1$ | Repeat transposition step.<br>Ensure cell viability as a<br>population of cells with highly<br>accessible DNS (eg.,<br>activated granulocytes, dead or<br>dying cells, unsupported<br>organisms). |
|  | NFR peak detected | No | Yes |  | Repeat library preparation.<br>Quantify library again to ensure<br>frequent size distribution.<br>Remove sources of sample<br>degradation. Repeat initial<br>sample preparation. |
|  | Mononucleosomal peak detected | No | Yes |  | Repeat initial sample<br>preparation. Quantify library<br>again to ensure fragment size<br>distribution. Check library has<br>200bp peak. |
| | TSS enrichment | <4 | [4, 6) | $\geq 6$ | Consider pre-treating cells with<br>DNase or using flow cytometry<br>to sort viable cells.<br>Low score may<br>indicate poor sample<br>preparation, poor sample<br>quality,<br>a population of cells with highly<br>accessible DNA (eg.,<br>activated granulocytes, dead or<br>dying cells, unsupported<br>organisms). |
| ChIPmentation | Uniquely mapped | <60% | [60% ,80%) | $\geq 80\%$ | Increase initial cell numbers. |
| | Sequence length | <50 bp | $\geq 50$ bp | | Remove sources of sample<br>degradation. |
| MethylationEPIC | % failed probes | >10% | [10%, 1%) | $\leq 1\%$ | Ensure optimal amount of input<br>DNA for the bisulfite<br>conversion kit.<br>Optimize PCR conditions in<br>whole genome isothermal<br>amplification. |
|  | Beta value distribution | >2 peaks | 2 peaks |  | Remove unreliable probes.<br>Remove sources of background<br>contamination. |
|  | %CpG methylation | <20% or >80% | [20%, 80%] |  | Repeat bisulfite conversion and whole genome amplification. |
| MeDIP-seq | Sequencing depth | <30M | $\geq 30M$ | | **<br>Repeat DNA extraction and purification. Repeat immunoprecipitation. |
| | CpG coverage | <40% | [40%, 60%) | $\geq 60\%$ | If caused by nonspecific binding, consider magnetic beads for immunoprecipitation instead of agarose beads. Siliconized tubes can also mitigate the non-specific binding of DNA to tube walls. Alter antibody and DNA incubation time. |
| | Sequence length | <50 | $\geq 50$ | | **<br>Repeat DNA extraction and purification. Repeat immunoprecipitation. |
| MINT-ChIP | Uniquely mapped | <60% | [60%, 80%) | $\geq 80\%$ | ** |
| | Uniquely mapped reads | <2M | [2M, 3M) | $\geq 3M$ | *<br>** |
| | Reads in peaks | <35 | [35, 50) | $\geq 50$ | Incorporate total Histone (H3) as control. |
| | Useful reads efficiency | <70% | $\geq 70\%$ | | Repeat IP reaction and amplification. |
| | PCR duplicates | $\geq 50\%$ | <50% | | Repeat IP reaction and amplification. |
| | Sequence length | <50 bp | $\geq 50$ bp | | * |
| miRNA-seq | Sequencing depth | <4M | [4M, 8M) | $\geq 8M$ | Increase starting material. Remove sources of RNA contamination. |
| | Aligned reads | <2M | $\geq 2M$ | | Repeat library construction. Ensure proper adaptor ligation in library construction. Use adapters with random sequences so that there is a greater probability of adapter ligating to a miRNA. |
| | Sequence length | <17 bp | $\geq 17$ bp | | Remove sources of RNA contamination. Repeat size exclusion steps. |
| Multiome | Number of cells | <500 | [500, 1000) | $\geq 1000$ | Check nuclei for proper quantification. Ensure proper nuclei handling. Remove background RNA and DNA. |
| | Median UMI per cell | <500 | [500, 1500) | $\geq 1500$ | Possible improper emulsification, partially or nonhomogeneous GEM emulsification reactions can occur which can be detected by visually inspecting pipette tips throughout steps. If this occurs, re-preparing the sample is the recommendation from 10x Genomics. |
| | Reads mapped to exons | <40% | [40%, 60%) | $\geq 60\%$ | *<br>** |
| | Transcriptomic reads in cell | <40% | [40%, 70%) | $\geq 70\%$ | *<br>**<br>Repeat sample preparation. Repeat library preparation. If transcriptomic reads low due to a population of nuclei with low RNA content, rerun pipeline with appropriate cell calling parameters. |
| | Cells with mitochondrial reads | <50% | [50%, 75%) | $\geq 75\%$ | Repeat tissue extraction avoiding necrotic cells. Repeat cell isolation. |
| | Mapped reads | <50% | [50%, 75%) | $\geq 75\%$ | Repeat library construction. |
| | Fragments in peaks | <5% | [5%, 10%) | $\geq 10\%$ | Ensure nuclei integrity.<br>Repeat transposition step.<br>Repeat emulsification.<br>Repeat sample extraction as low values could be from population of cells with un-compacted DNA (activated granulocytes, dead or dying cells.) |
| | GEX sequence length | <50bp | $\geq 50$bp | | *<br>**<br>Repeat library preparation. |
| | ATAC sequence length | <37bp | $\geq 37$bp | | *<br>**<br>Repeat library preparation. |
| RNA-seq | Sequencing depth | <20M | [20M, 50M) | $\geq 50$M | Increase sample input for sequencing. |
|  | %GC | <20% or >80% | [20%, 40%) or (60%, 80%] | [40%, 60%] | Optimize PCR parameters for cDNA library construction. |
| | %rRNA | >20% | (10%, 20%] | $\leq 10\%$ | Enrichment for mRNA with polyA selection.<br>Consider rRNA depletion methods. |
| | %globinRNA | >30% | (15%, 30%] | $\leq 15\%$ | Consider globin depletion methods.<br>Repeat PBMC extraction ensuring exclusion of RBCs. |
|  | mRNA reads | <16M | [16M, 30M) | ≥30M | Enrichment for mRNA with polyA selection methods.<br>Consider rRNA removal kit.<br>Consider RRNA removal kit. |
|  | %uniquely mapped | <40% | [40%, 75%) | ≥75% | Increase library diversity by increasing/improving quality RNA input for cDNA library.<br>Optimize PCR parameters for cDNA library construction.<br>** |
|  | Sequence length | <50 bp | ≥50 bp |  | *<br>** |
| scATAC-seq | Mapped reads | <50% | [50%, 75%) | ≥ 75% | Repeat library construction. |
|  | Fragments in peaks | <5% | [5%, 10%) | ≥10% | Repeat transposition step.<br>Repeat sample extraction as low values could be from population of cells with un-compacted DNA (activated granulocytes, dead or dying cells).<br>Ensure nuclei integrity. |
|  | Sequence length | <37 bp | ≥37 bp |  | *<br>**<br>Repeat library preparation. |
| scRNA-seq | Number of cells | <500 | [500, 1000) | ≥1000 | Repeat cell preparation.<br>Optimize cell preservation methods.<br>Repeat library construction. |
|  | Median UMI per cell | <500 | [500, 1500) | ≥ 1500 | Increase input material for cDNA synthesis.<br>If sample quality is strong, increasing the PCR cycles by two extra cycles during cDNA amplification can bring transcripts above background levels. |
|  | Reads mapped to exons | <40% | [40%, 60%) | ≥60% | *<br>** |
|  | Transcriptomic reads in cell | <40% | [40%, 70%) | ≥70% | *<br>** |
|  | Cells with mitochondrial reads | >50% | (20%,50%] | ≤20% | Repeat tissue extraction avoiding necrotic cells. Repeat cell isolation. Implement measures to ensure cellular quality and prevent cell lysis. |
|  | Sequence length | <50 bp | ≥50 bp |  | *<br>** |
| snmC-seq | % Genome | <1% | [1%, 5%) | ≥5% | *<br>** |
|  | CCC methylation rate | >3% | (2%, 3%] | ≤2% | Repeat bulsulfite conversion and subsequent steps. |
| | %uniquely mapped | <40% | [40%, 70%) | $\geq 70$ | Increase input material. |
| | Reads per cell | <10K | [10K, 1M) | $\geq 1M$ | *<br>** |
| | Cell level mCG rate | <0.5 | [0.5, 0.7) | $\geq 0.7$ | Repeat bisulfite conversion and subsequent steps. |
| | Cell level mCH rate | $\geq 0.08$ | <0.08 | | Repeat bisulfite conversion and subsequent steps. |
| | Sequence length | <30 bp | $\geq 30$ bp | | ** |

Software components for the assessed and containerized pipelines are included for each pipeline in Table 10. To determine QC metrics, the pipelines utilize fifteen open source tools as well as two internally developed programs: *SeqLengths* which characterizes the average FASTQ read sequence length, and *QC Parser*, which generates individual JSON files for each sample’s QC metrics. The scATAC, scR-NAseq and multiome pipelines also utilize the 10X Genomics pipelines which can further streamline downstream analysis.

**Table 10.**
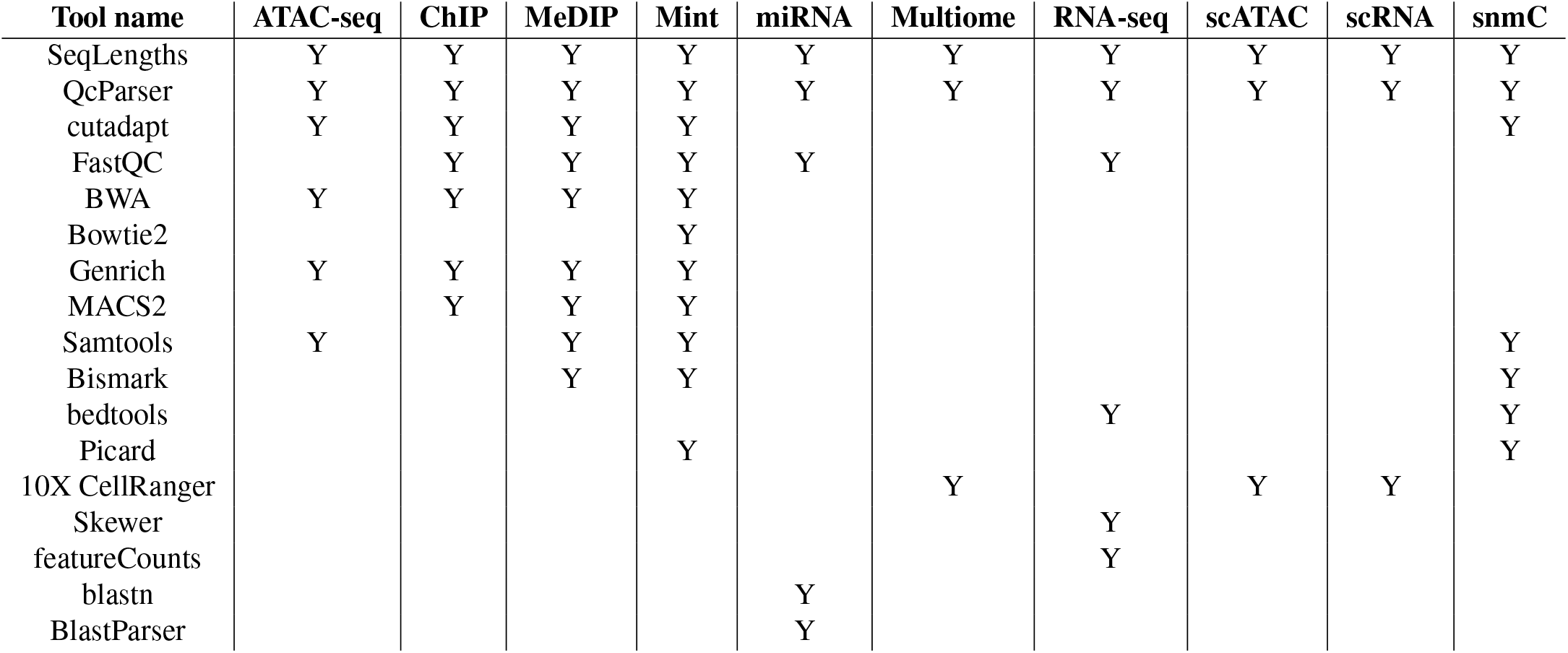
Pipeline components.

| Tool name | ATAC-seq | ChIP | MeDIP | Mint | miRNA | Multitome | RNA-seq | scATAC | scRNA | snmC |
| --- | --- | --- | --- | --- | --- | --- | --- | --- | --- | --- |
| SeqLengths | Y | Y | Y | Y | Y | Y | Y | Y | Y | Y |
| QcParser | Y | Y | Y | Y | Y | Y | Y | Y | Y | Y |
| cutadapt | Y | Y | Y | Y |  |  |  |  |  | Y |
| FastQC |  | Y | Y | Y | Y |  | Y |  |  |  |
| BWA | Y | Y | Y | Y |  |  |  |  |  |  |
| Bowtie2 |  |  |  | Y |  |  |  |  |  |  |
| Genrich | Y | Y | Y | Y |  |  |  |  |  |  |
| MACS2 |  | Y | Y | Y |  |  |  |  |  |  |
| Samtools | Y |  | Y | Y |  |  |  |  |  | Y |
| Bismark |  |  | Y | Y |  |  |  |  |  | Y |
| bedtools |  |  |  |  |  |  | Y |  |  | Y |
| Picard |  |  |  | Y |  |  |  |  |  | Y |
| 10X CellRanger |  |  |  |  |  | Y |  | Y | Y |  |
| Skewer |  |  |  |  |  |  | Y |  |  |  |
| featureCounts |  |  |  |  |  |  | Y |  |  |  |
| blastn |  |  |  |  | Y |  |  |  |  |  |
| BlastParser |  |  |  |  | Y |  |  |  |  |  |

### microRNA-seq

microRNA sequencing (microRNA-seq) sequences miRNA, a type of RNA that is 21 – 24 nt long that regulates gene expression at the transcriptional or post-transcriptional level by binding to complementary mRNAs. MicroRNAs are expressed in the nucleus as primary-miRNA (pri-miRNA), processed by the Drosha enzyme into precursor-miRNA (pre-miRNA), then exported to the cytoplasm where they are further cleaved by the Dicer enzyme into 21-24 nt double stranded mature miRNA. miRNAs then bind to Argonaute effector proteins for their downstream regulatory function. Abnormal miRNA expression has been implicated in many human diseases, including infectious disease. (67, 68) Due to their conserved function in gene expression, an understanding of the changes in miRNA expression between healthy and disease states or before and after an infection or exposure to chemicals can provide insights into mechanistic pathways, novel biomarkers, and enable development of predictive signatures of infection or exposure.

Pre-processing steps for samples include sample acquisition, total RNA extraction (Norgen total RNA extraction kit), enrichment of sRNA and enzymatic modifications. There are several methods that can be used for extraction of total RNA and enrichment of sRNA, each of which have potential to introduce bias. Due to the high abundance of other RNA species in total RNA, most of which are greater in length than miRNA, it is important to maintain the integrity of the total RNA to avoid degradation into smaller length RNA which could confound extraction of miRNA based on their size range. A total size distribution from the read alignment should show distinct peaks in the 21-24 nt range which would otherwise indicate problems with the input material or library preparation. In addition, mapping of reads to known miRNA loci can be done using a database such as miRBase to ensure majority of the reads are in known miRNA genomic regions. Based on these considerations, sequencing length should be greater than 17bp. Sequences under 17bp may be indicative of degradation and sequences over this may require repeating the size exclusion steps.

Libraries were prepared using the QIAseq miRNAseq Library kit with 100 ng total RNA input. miRNAs can have modifications at their 5’ and 3’ ends which require appropriate enzymatic modifications to enable proper adapter ligation during the library preparation step. If proper enzymatic modifications are not done it can introduce errors. Once the samples are ready for library preparation, one of several ways to prepare a library can be used, each of which could have their own specific biases. In general, the steps in library preparation include possible polyA tailing, ligation of sequencing adapter, PCR primer and sample barcode for multiplexing, cDNA synthesis by reverse transcription and PCR amplification. Improper adapter ligation is usually the greatest source of bias which can lower the percentage of aligned reads and misidentification of differentially expressed miRNAs. One possible solution is to use adapters with random sequences so that there is a greater probability of adapter ligating to a miRNA. In this strategy all the different adapter sequences should be trimmed in the downstream bioinformatics analysis steps before proceeding with the data analysis. Sequencing was performed using the Illumina NextSeq 500 High-Output 74bp single-end read with 5-10M read clusters per sample. Optimal samples will have a sequencing depth between 4M and 8M with aligned reads over 2M.

The primary cause of higher PCR duplication rate is low starting material or mistakes in the steps preceding the library preparation. High PCR duplication can result in libraries with low complexity which can lead to missing miRNA species in the data analysis step and lowering the percentage of reads mapped and aligned. Input sample quantity and quality in addition to the pre-processing steps should be optimized for minimizing PCR duplication rate.

### snmC-seq

Single-nucleus methylcytosine sequencing (snmC-seq) characterizes nucleotide modifications at a single-nucleosome resolution level by measuring the occurrence of 5-methyl- and 5-hydroxymethylcytosines. Profiling single cell methylomes enables cell state delineation and provides insight to the cell cycle stage, transcriptional activity, mitotic age and proliferation potential (69).(40). This single-cell epigenomics assay has been used to identify and characterize diverse neuronal types and offers scalability to characterize cellular populations traditionally difficult to identify (34).

We isolated populations of nuclei by fluorescence-activated cell sorting (FACS) that were highly enriched for neurons (NeuN+) or glia (NeuN-) from human and mouse adult frontal cortex tissue. (70). Isolated nuclei were labeled with AlexaFluor488 conjugated anti-NeuN antibody (MAB377X, Millipore) and sorted by flow cytometry. Single nucleus methylome library preparation began with bisulfite conversion of single nuclei (Zymo EZ-96 DNA Methylation-Direct TM Kit). Four indexed random primers were used to synthesis bisulfite converted single nuclei in a 96 well plate. Samples were bead purified (DynaMag™-96 Side Magnet, ThermoFisher). Samples were pooled for an adaptase reaction to attach adapters for sequencing. (Accel-NGS Adaptase Module for Single Cell Methyl-Seq Library Preparation, Swift Biosciences). Samples were sequenced on an Illumina HiSeq 4000.

Optimal samples provide at least 1% of genome coverage. The methylation rate of the trinucleotide sequence CCC is assessed as a metric as an estimate for the rate of bisulfite non-conversion. Low input material can be the cause of low and failed uniquely mapped reads metric, specifically, the Smart-Seq2 protocol purification step can lead to a significant loss of material. Methylation of CG sites (mCG) rates andmethylation of non-CG (mCH) rates are metrics indicative of both quality and accuracy of the assay. mCG and mCH rates should reflect a genomic methylation state: high mCG occurance as 60% -90%of all CpG sites in mammals are methylates and very low mCH rates as methylation outside of CpG sites are rare. (71) Repeating bisulfite conversion and subsequent steps would be necessary if mCG and mCH rates were significantly out of passing range.

### MeDIP-seq

Methylated DNA immunoprecipitation sequencing (MeDIP-seq) couples a magnetic bead-based antibody capturing method with NGS to agnostically assess genomic methylation. MeDIP-seq assesses genomic methylation by characterizing the modifications of cytosines: 5mc (5-methylcytosine) or 5hmc (5-hydroxymethylcytosine). MeDIP-seq has been used to demonstrate environmentally induced epigenetic transgenerational inheritance of disease and phenotypic variation. (72)

Genomic DNA from PBMCs was extracted and purified (Promega Genomic DNA Purification Kit) followed by sonication (Covaris 300bp protocol). DNA was heat denatured into single stranded DNA to improve antibody binding affinity. DNA was incubated with anti-5mC (5-methylcytosine) antibodies (monoclonal mouse anti 5-methyl cytidine; Diagenode #C15200006). The DNA-antibody complex was captured by magnetic beads (Dynabeads M-280 Sheep antiMouse IgG; 11201 D) and unbound DNA was removed. Samples were treated with proteinase K to digest the antibody and eluted DNA was prepared for NGS (NEBNextVR UltraTM RNA Library Prep Kit for Illumina).

An optimized immunoprecipiation reaction with high quality DNA input is required to achieve passing metrics. CpG coverage percent can be affected from non-specific binding in immunoprecipitation. Because antibody-based selection is biased toward hypermethylated regions, screening could exclude CpG islands if they are hypomethylated. Poor CpG coverage can be caused from nonspecific binding. Antibody and DNA incubation time can also alter binding results which can influence sequenced motifs.

### ChIPmentation

Chromatin Immunoprecipitation with on bead tagmentation (ChIPmentation) introduces an expedited library preparation method for ChIP-seq. In tagmentation a hyperactive Tn5 transposase simultaneously fragments DNA and ligates adaptors directly to bead bound immunoprecipitated chromatin. In combination with next generation sequencing, chIpmentation maps histones and their post translational modifications of transcription factors throughout the genome. This method offers efficiency, reduces cost and greatly reduces input requirements (73).

sequencing library preparation by Tn5 transposase (Nextera DNA Sample Prep Kit Illumina and MinElute Kit). Experimental datasets can vary from high to low quality. Several assays (ATAC-seq, ChIPmentation, MeDIP-seq, Mint-ChIPseq, RNA-seq, and others) rely upon amplification of target sequences with PCR. When source target nucleic acids are limited or experimental techniques are in need of improvement, excessive amplification of duplicate target sequences can result. Early identification of low quality datasets enables 1) avoidance of low quality data into downstream analytics, 2) enables monitoring of experimental assays for early identification of possible issues, and 3) enables assay selections samples with limited cells or DNA available.

ChIPmentation assays can suffer from high PCR duplication rate due to the low immunoprecipitated DNA yield that requires a high number of PCR amplification cycles to generate a sufficient library for sequencing. This decreased complexity of input material for sequencing can cause high PCR duplication rates. Bias can also arise from Tn5 insertion frequency being higher in positions surrounding the center of nucleosomes where the DNA is most accessible (73). Samples should have a uniquely mapped reads percentage over 80%. Increasing initial cell numbers can increase the number of mapped reads (36). Optimal chIPmentation samples should have a sequence length of over 50 bp. Any sample degradation leading to protein degradation will innately lead to poor sample sequencing as chromatin proteins are protecting the DNA during the tagmentation process and are the link to accurately relate the DNA sequence to the antibody binding to the proteins around it. Increasing initial cell numbers can increase the number of mapped reads (36).

## Discussion

Outlining high quality assay metrics for 11 different epigenomics and transcriptomics datasets alongside mitigation strategies to assist in optimizing protocols to yield high quality results will alleviate the issues in complex datasets that are difficult to navigate due to dynamic research teams that exhibit a multitude of research objectives. On projects where there may be numerous different handlers of a sample from collection to sequencing, it can be difficult to pinpoint the source of a sample’s problem or contamination. QC should be a prioritized step in all computational epigenomic and transcriptomic workflows. Although many workflows incorporate QC steps, the advantages of looking at individual sample quality within a given datasets is often overlooked. The ability to assess datasets for QC issues at the sample level prior to downstream analysis prevents the misinterpretation of results and serves as a strong indicator of bench protocols that yield high quality and reproducible results. Eliminating analysis on low quality data can optimize time and reduce the probability of developing erroneous results.

The assessed pipelines were containerized in Singularity and designed for a high-performance computing (HPC) environment which offers great advantages in computing speed and reproducibility. The HPC environment is well suited for the QC of epigenetic and transcriptomic datasets because it enables the parallelization of pipeline steps to achieve rapid QC metric reporting. Singularity also offers the ability to run containers in internet starved environments and on a shared environment without root access. Packaging software offers large research teams from different institutions and projects that span a long period of time to achieve reproducible results. All of these advantages offer great interoperability for research teams involving multiple collaborators and reproducibility of results to ensure datasets from different time points are being analyzed with the identical tools, algorithms, and standardized set of metrics.

## Summary

This paper serves a comprehensive review and guide for epigenomics and transcriptomics assay quality. The assessed QC pipelines as containerized open source code to be used on high-performance computing environments are available as an open-source package (https://github.com/mitll/Omics_QC_pipelines).

## Acknowledgements

Funding and guidance were provided by the Defense Advanced Research Projects Agency; Epigenetic CHaracterization and Observation (ECHO) program. The authors acknowledge the MIT SuperCloud and Lincoln Laboratory Supercomputing Center for providing (HPC, database, consultation) resources that have contributed to the research results reported within this paper, Darrell Ricke, PhD, Philip D. Fremont-Smith, and Catherine Cabrera, PhD (MIT LL), Joseph Ecker (Salk Institute) and Thomas Thomou (DARPA).

## Legal and funding statement

DISTRIBUTION STATEMENT A. Approved for public release. Distribution is unlimited.

This material is based upon work supported by the Defense Advanced Research Projects Agency under Air Force Contract No. (FA8702-15-D-0001). Any opinions, findings and conclusions or recommendations expressed in this material are those of the author(s) and do not necessarily reflect the views of the Defense Advanced Research Projects Agency.

## Footnotes

1 Note: There are many other QC tools for alignments, but the included examples are meant to showcase key QC metrics of interest rather than the specific functionality of Picard itself.

